# Bulk Transcriptomic Analysis with InMoose, the Integrated Multi-Omic Open-Source Environment in Python

**DOI:** 10.1101/2024.11.29.625982

**Authors:** Maximilien Colange, Guillaume Appé, Léa Meunier, Solène Weill, W. Evan Johnson, Akpéli Nordor, Abdelkader Behdenna

**Affiliations:** Epigene Labs, Paris, France; Rutgers New Jersey Medical School, Rutgers University, Newark, NJ, USA

## Abstract

We introduce InMoose, an open-source Python environment aimed at omic data analysis. We illustrate its capabilities for bulk transcriptomic data analysis.

Due to its wide adoption, Python has grown as a *de facto* standard in fields increasingly important for bioinformatic pipelines, such as data science, machine learning, or artificial intelligence (AI). As a general-purpose language, Python is also recognized for its versatility and scalability. InMoose aims at bringing state-of-the-art tools, historically written in R, to the Python ecosystem. Our intent is to provide a drop-in replacement for R tools, so our approach focuses on the faithfulness to the original tools outcomes. The first development phase has focused on bulk transcriptomic data, with current capabilities encompassing data simulation, batch effect correction, and differential analysis and meta-analysis.

## Introduction

Since the first human genome was sequenced in 2000, omic profiling technologies have seen their costs reduced by multiple orders of magnitude, and omic profiling is now performed routinely. Petabytes of omic data are generated each year, yet most of it remains largely under-used.

Bridging the gap between data science, informatics and biology remains a key challenge to enable the analysis of such massive and complex data. Overcoming this challenge requires the development of efficient, easy-to-use, and interoperable tools, to let practitioners focus on scientific questions rather than technical issues.

Despite the historical prevalence of R for data analysis and visualization applications, Python is gaining momentum in the bioinformatics and biostatistics landscape. As signs of the trend, we note that Python has maintained a higher popularity than R in the past decade (1), and that more and more tools are developed directly in Python (such as *lifelines*, a library dedicated to survival analysis (2), or *scanpy*, a library dedicated to single-cell RNA-Seq data analysis (3)). We have now reached a point where reference tools may be only available in Python, such as for RNA velocity and trajectory analysis. Compatibility with Python has therefore become more crucial than ever.

We detail the reasons why we believe this trend is likely to amplify in the foreseeable future. First and foremost, Python has become a language of choice in data science, machine learning and AI, which are becoming integral parts of modern bioinformatic pipelines. As a general-purpose language, Python is more versatile than R, geared towards statistical manipulations and data visualizations. For instance, Python has always been a more powerful tool than R for managing and manipulating strings (*e*.*g*. DNA sequences). Python’s overall ecosystem facilitates the integration of bioinformatics tools into large-scale frameworks (*e*.*g*. web-based interactive interface), increasing their functionality and widening their targeted audience. Finally, Python is a common target language for data management infrastructure API (cloud providers, public databases…): this makes such integrations smoother as Python can be used both for tool development, data management and platform deployment.

These advantages motivate the further development of the Python bioinformatics ecosystem. We advocate the importance of three key factors:

- providing a single-language unified ecosystem for data acquisition, management, analysis, visualization, and democratization, in the hope to eliminate the need to interoperate several languages (typically R and Python). Cross-language interfaces are a common source of technical difficulties and of computational inefficiency, pain points that grow harder as the amount of data grows.
- ensuring retro-compatibility (through reproducibility of results) with the large R-based legacy. Transitioning to a new ecosystem makes little sense if all previous results need to be re-computed. To that end, tools from both ecosystems should have similar if not identical results.
- maintaining open tools to ease dissemination and collaboration. As trans-disciplinary work becomes increasingly important in modern bioinformatics, this element drives our open-source approach. We hope to leverage Python’s popularity to invite developers, scientists and engineers from other disciplines to learn about and ultimately contribute to the bioinformatics ecosystem.

To address those two points, we introduce InMoose (Integrated Multi-Omics Open Source Environment), an open source Python unified framework for -omic data type. The first development phase of InMoose focuses on bringing to the Python world tools for the manipulation of bulk transcriptomics data (microarray, RNA-Seq). It features data simulation, data clustering, and batch effect correction as well as differential expression analysis, and meta-analysis, for microarray and RNA-Seq data analysis. InMoose is accessible at https://github.com/epigenelabs/inmoose and is released under a GNU GPL3 license.

InMoose demonstrates the advantages of our approach:

- it is developed with the intent to capitalize on and facilitate interactions with other popular Python data-oriented libraries (such as *numpy, scipy* or *pandas*)
- it focuses on reproducing in Python the results obtained in R, so as to ensure the compatibility of both ecosystems (4)
- it demonstrates significant computational performance improvements (4)
- our package integrates easily into web-based user-friendly analyses platforms (*e*.*g*. Epigene Labs’ proprietary mCUBE platform)
- it provides an opportunity window to improve both functionality and performance while porting existing code from one language to another

Prominent features of InMoose are based on state-of-the-art tools: *splatter* (5) (data simulation), *consensus clustering* (6) (cohort stratification), *ComBat* (7) and *ComBat-Seq* (8) (batch effect correction), *limma* (9), *edgeR* (10), *DESeq2* (11) (differential expression analysis). Because implementation details do matter, in most cases we have chosen to port R tools to Python, following the original source code as faithfully as possible, instead of re-implementing them. Only *ComBat* (batch effect correction for microarray data) and *splatter* (RNA-Seq and single-cell RNA-Seq data simulation) were re-implemented, at a time where our porting approach was not completely defined. Note that the behavior of InMoose against the original R *ComBat* was assessed in a dedicated publication (4). As for *splatter*, our re-implementation is based on the original article describing the simulation model (5) and on the original source code to clarify specific points.

InMoose also offers original features:

- a cohort quality control report, which gathers various metrics to assess the quality of an aggregated cohort after batch effect correction; and
- a differential expression meta-analysis tool, aimed at aggregating differential expression analysis results obtained on different cohorts and/or from different tools.

## Results

We illustrate the range of capabilities of InMoose through an example workflow of differential expression meta-analysis on simulated RNA-Seq data. This provides concrete code showing how to use and integrate InMoose in real-life workflows.

The example workflow starts with the simulation of a RNA-Seq cohort, featuring 3 batches and 2 groups. It then proceeds to two differential expression meta-analyses:

- one by first running differential expression analyses on each batch then aggregating the results with a random-effect model. We call it the AD (Aggregate Data) approach.
- one by first aggregating the batches into a single cohort, correcting batch effects then running a differential expression analysis on the resulting cohort. We call it the Individual Sample Data (ISD) approach.

The overall workflows are illustrated on Figure 1. The AD/ISD terms are inspired by terminology used in clinical trials meta-analysis (12).

**Figure 1.**
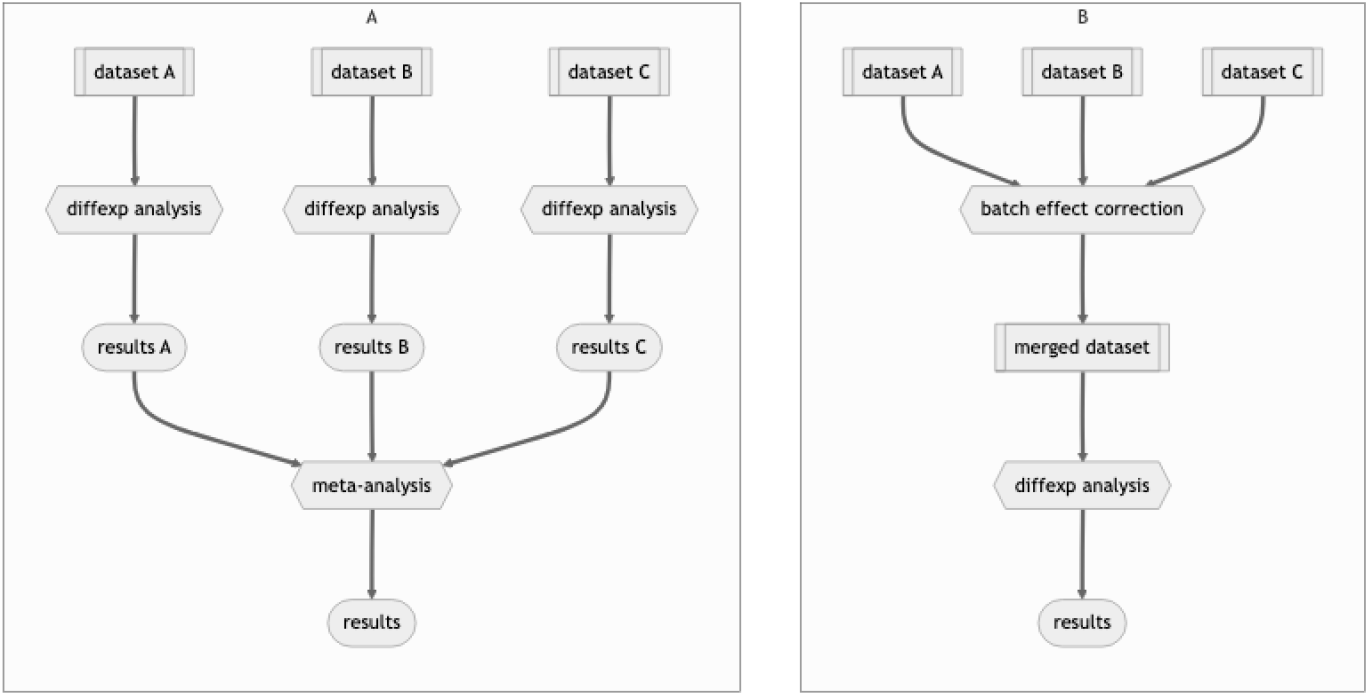
Conceptual illustration of the meta-analysis workflows. **A**. Aggregate Data (AD) workflow. **B**. Individual Sample Data (ISD) workflow.

The code to simulate RNA-Seq data is presented in Listing 1. Lines 1 to 3 import packages and functions used in the remainder of the code. Lines 5 through 27 define the parameters for the simulation as well as a few helper variables. Data simulation is performed on line 32.

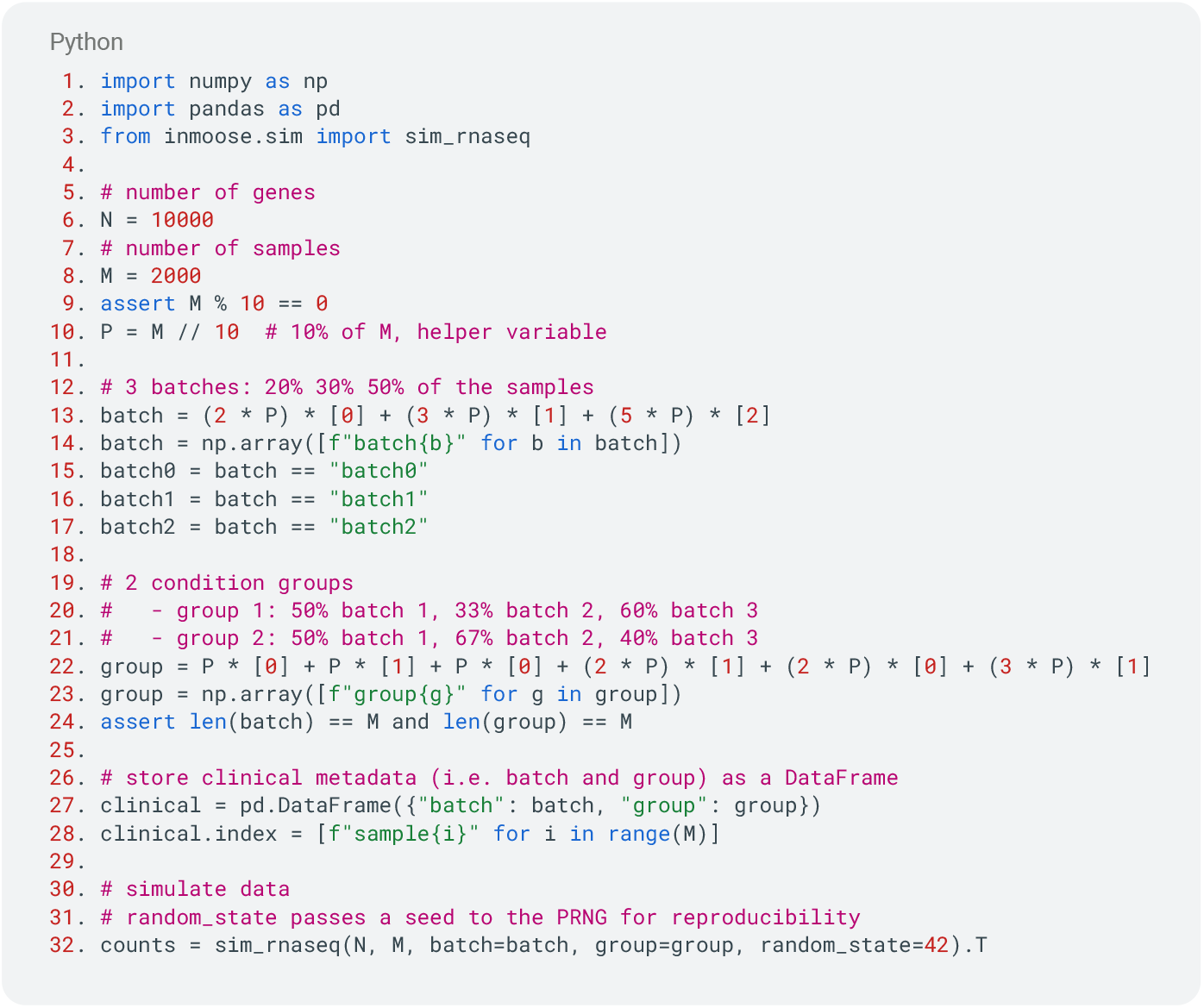

**Listing 1.** Code for simulating RNA-Seq data.

The code for the meta-analysis workflows is presented in Listing 2. It assumes the molecular data is stored in a variable *counts*, and the clinical metadata is stored in a variable *clinical* as a pandas DataFrame, as done in Listing 1. It also assumes that variables *batch0, batch1* and *batch2* are mask index corresponding to each individual batch – see Listing 1, lines 15 through 17. Lines 1 to 3 import packages and functions used in the remainder of the code. AD meta-analysis occurs on lines 5–11. ISD meta-analysis occurs on lines 13–18.

The code of Listing 1 and 2 is intended to run with version 0.7.3 of InMoose, and does not require any other dependencies – *numpy* and *pandas* are dependencies of InMoose and are automatically installed when installing InMoose. It does not require any specific hardware and was run in 3 minutes on a laptop (2022 MacBook Pro).

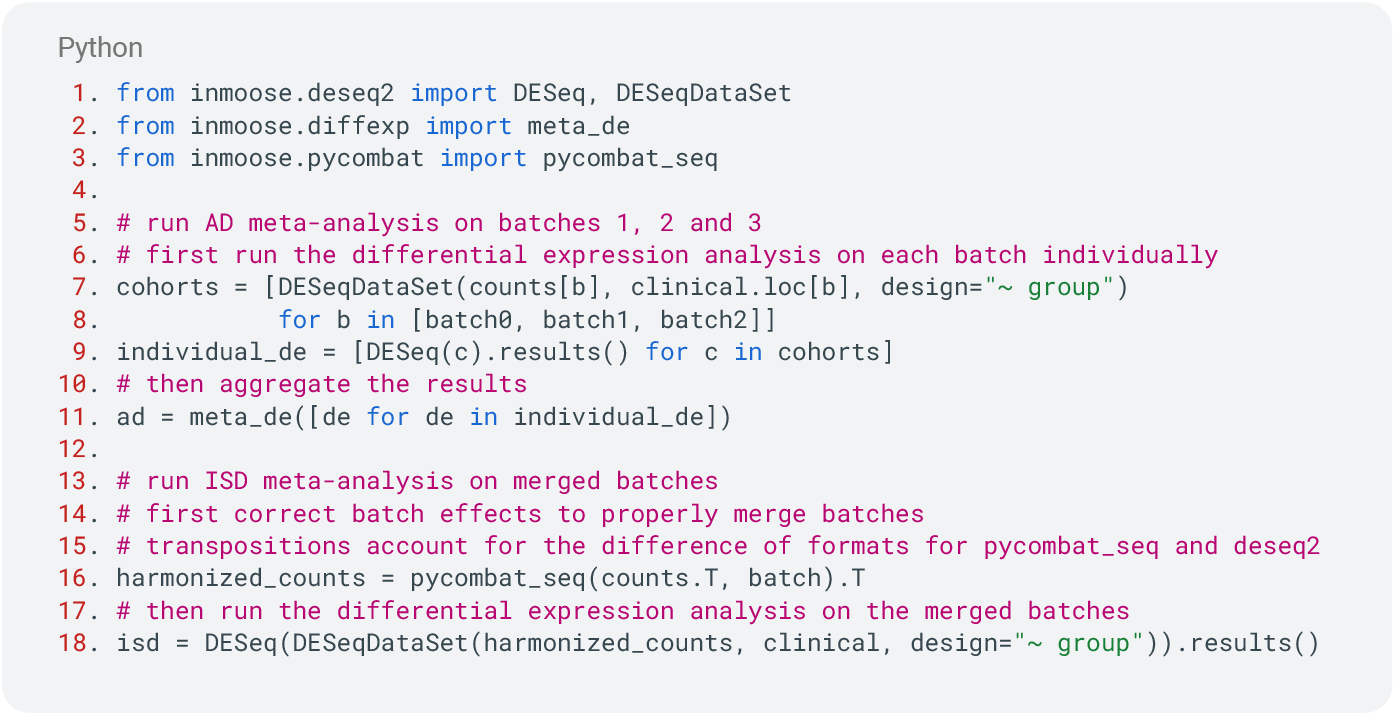

**Listing 2.** AD and ISD meta-analysis code.

Beyond the mere correction of batch effects to integrate data, InMoose can generate a HTML report assessing the quality of the cohort. This report gathers several metrics and diagrams, evaluating for instance the quality of the correction of batch effects, or the correlation of data with the covariates. An example of cohort QC report is featured in the annex.

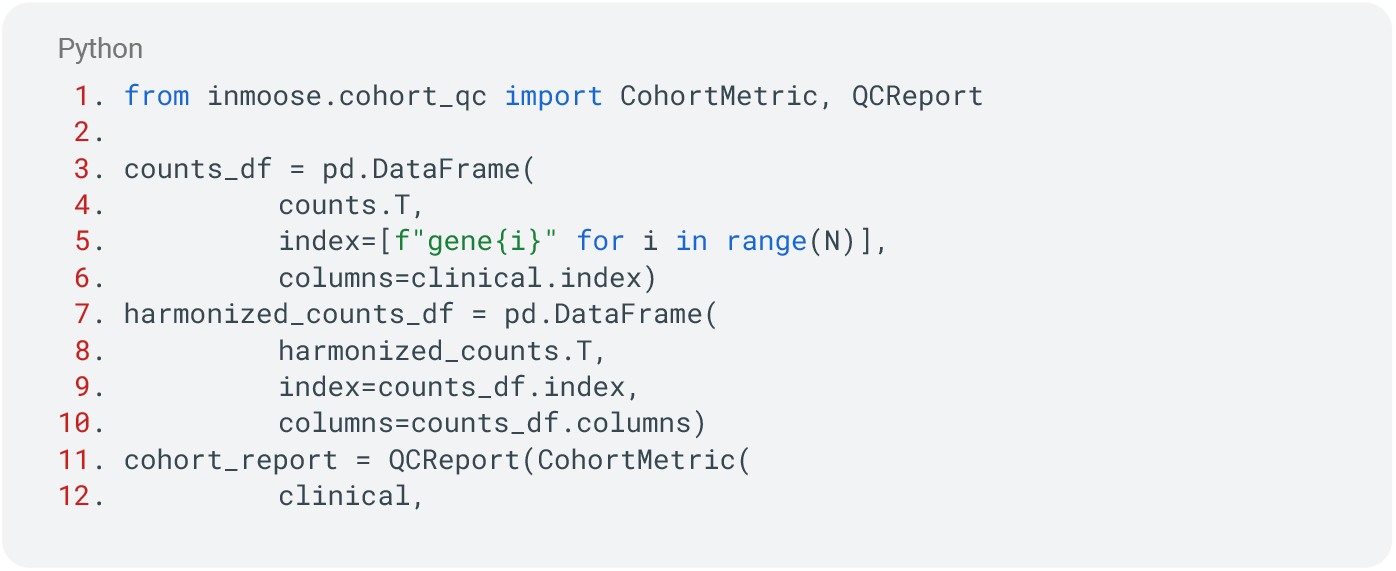
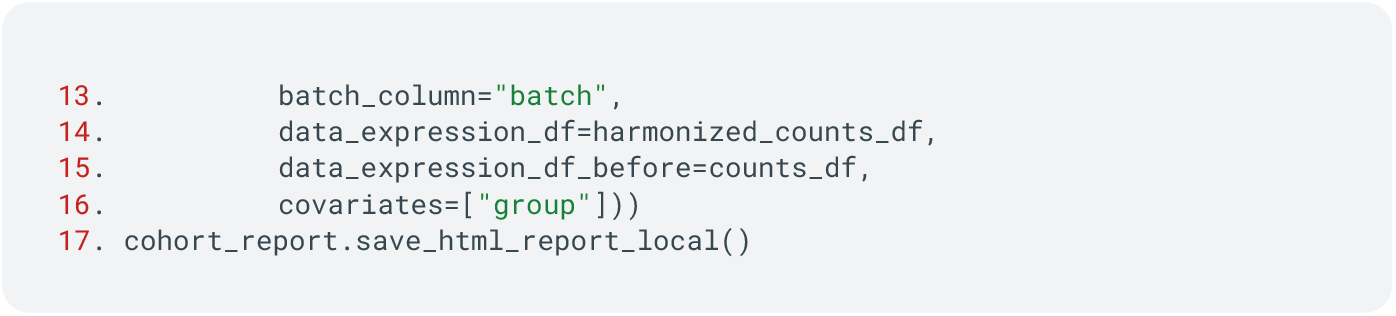

**Listing 3.** Code snippet to generate cohort QC report.

We now comment on the results obtained by this workflow to illustrate the capabilities of InMoose. We remind the reader that they are applied to simulated data and that the behavior on real-life data may differ.

Figures 2A and 2B show the whole cohort plotted in PCA space, respectively before and after batch effect correction. Data points are colored by batch and by group. We clearly see 6 clusters before batch effect correction, corresponding to all 6 batch-group combinations. Those are reduced to 2 clusters after batch effect correction, corresponding to the two groups. These figures highlight the quality of the batch effect correction by *pycombat*. For a thorough evaluation of the abilities of *pycombat*, we refer the reader to the dedicated publication (4).

**Figure 2.**
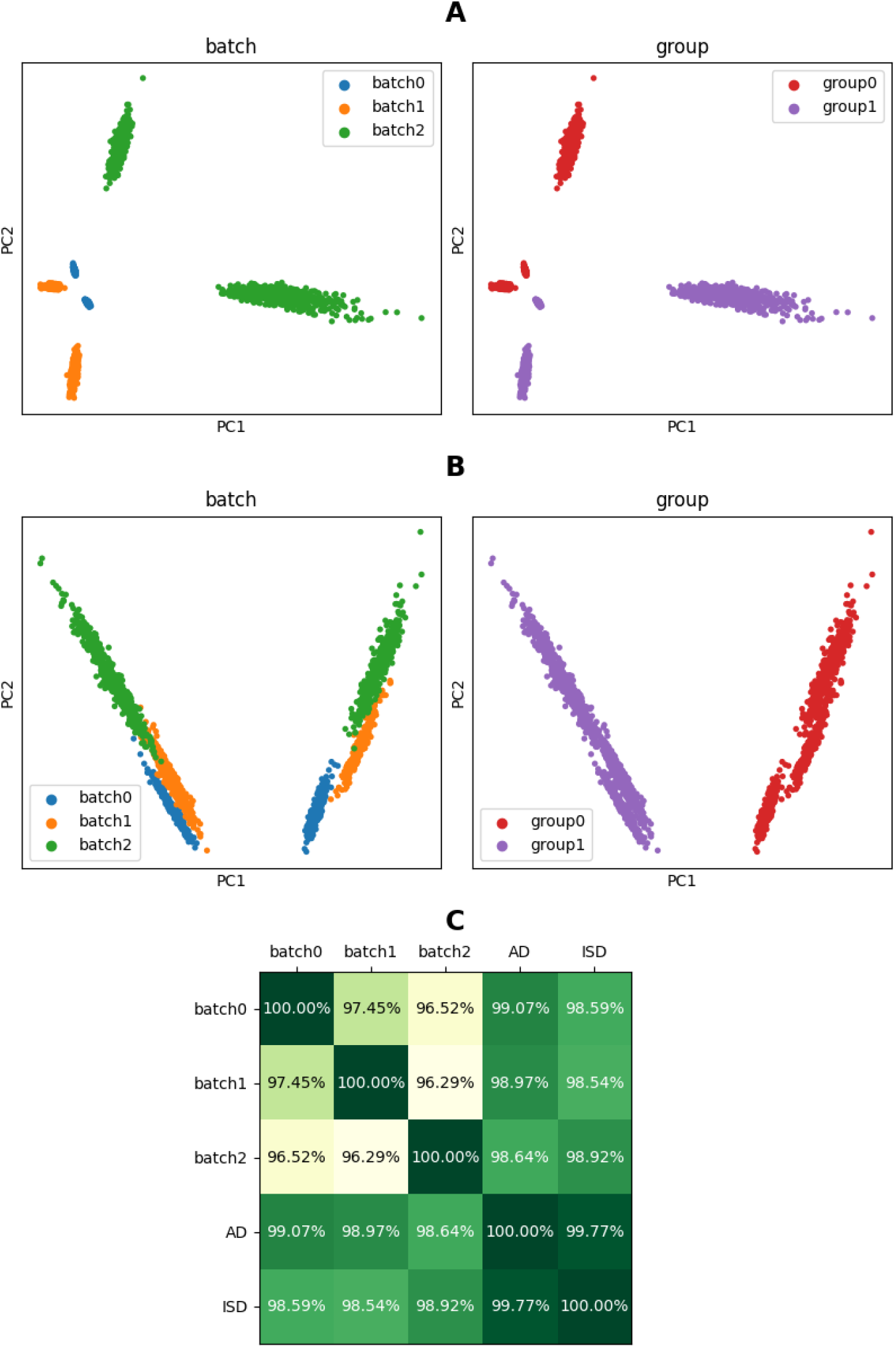
**A**. PCA plot of the cohort before batch effect correction. **B**. PCA plot of the cohort after batch effect correction. **C**. Pairwise log-fold-change correlations.

Figure 2C compares pairwise the log-fold-changes found in each individual cohort and in each meta-analysis (AD and ISD), as Pearson correlation values. The high correlations between log-fold-changes found in individual datasets indicate that DESeq2 accurately estimates the inter-group fold-change despite the presence of batch effects.

## Discussion

The bioinformatic landscape is mostly split between Python and R languages. For a given language, interoperability between packages is not immediate: it requires an effort to develop and adopt common data structures and/or data formats. In R, the Bioconductor ecosystem (13) achieves a high level of interoperability thanks to common data structures (such as *SummarizedExperiment* for gene expression data). It also fosters usability by providing a package management system and by requiring a standardized form of documentation as vignettes. On the Python side, a few core tools (such as PyPI for package management or readthedocs.org for documentation) can be considered as *de facto* standards given their wide adoption. On the bioinformatics part, the projects closest to Bioconductor might be *BioPython* (14), the *scverse* (15), as well as *omicverse* (16). All revolve around common data structures (Seq and SeqRecord for *BioPython*, AnnData and MuData for *scverse* and *omicverse*). They have grown a collection of Python packages around those common data structures, and follow a curation process to ensure that newly accepted packages are compatible with those data structures. Additionally, they promote common practices in terms of repository structure and of documentation. In this landscape, only *omicverse* and InMoose offer a single entry-point library instead of a collection of tools. The features of InMoose do not overlap significantly with those of *BioPython* or *scverse*. Of note, *omicverse* (as of version 1.6.7) wraps the batch effect correction features from an older version of InMoose, as well as the differential expression features from *pydeseq2*.

Interoperability between R and Python is also a growing concern. Bridging tools, such as the *rpy2* library (17) allows one to seamlessly call one language from the other. However, the need to translate and copy data structures from one language to the other, is a source of inefficiency, especially when the analysis goes back and forth between the two. We observe that Python is gaining popularity and momentum, as is exemplified by the scverse environment, *scanpy* for single-cell RNA-Seq data, or *pydeseq2* a reimplementation of DESeq2 R package into Python. Similarly, the initial features of InMoose are also reimplementations or ports of R packages.

We have attached importance to making the software open source, coupled with comprehensive documentation. We built a robust set of test cases, in an effort to encourage larger participation from the community. We believe that this will be benefiting the Python bioinformatics community and opening the way towards the translation of other widely used software from R to Python.

## Methods

InMoose ports to Python prominent features from well-established R packages, to provide a single entry point for bulk transcriptomic data analysis. Our approach is to be as faithful to the original tools as possible, as demonstrated by our studies (4,18), so as to provide a drop-in replacement tool for bioinformaticians willing to migrate their workflows from R to Python.

InMoose also features original capabilities to top those. In this section we detail the functionalities offered in InMoose, broken down by sub-module.

### a. pycombat – Batch Effect Correction

Batch effects are the product of technical biases, such as variations in the experimental design, technology platform, or even atmospheric conditions (19,20). They particularly reveal themselves when merging different datasets, which have likely been built under different conditions. If not corrected, these batch effects may lead to incorrect biological insight, since batch bias can mask signal of interest or can be wrongly interpreted as the product of a biological process.

The most commonly used methods for batch effects correction on high-throughput molecular data are *ComBat* (7) and *ComBat-Seq* (8), both implemented in the R library *sva* (21). They leverage an empirical Bayes approach for correcting the batch effect in, respectively, microarray and RNA-Seq datasets. Their similar mathematical frameworks robustly handle key limit cases, such as small sample sizes or the presence of outliers.

The *pycombat* module of InMoose features a Python port of *ComBat* and *ComBat-Seq*. It was shown to yield similar results as the reference R implementations in shorter computation times. While *ComBat* has another Python implementation (3), InMoose is, to our knowledge, the sole Python implementation of *ComBat-Seq*.

### b. edgepy, DESeq2, limma – Differential Expression Analysis

Transcriptomic data is often used as a proxy for protein expression, that drives the biological activity. Differential expression analysis aims at identifying genes whose expression level significantly differs from one group of samples to another. The difference in expression may be normal (*e*.*g*. when comparing different tissues) or pathological (*e*.*g*. when comparing healthy organ tissue and tumor tissue). In any case, differential expression analysis selects genes of interest to consider for *e*.*g*. mechanistic studies, biomarker identification or target-based therapeutic research.

*limma* (9) (for microarray data), *DESeq2* (11) and *edgeR* (10) (for RNA-Seq data) are state-of-the-art tools for differential expression analysis of bulk transcriptomic data. They all rely on a Bayesian approach that empirically estimates the contribution of each variable of interest (*e*.*g*. age, gender, condition, biopsy site…) to the gene expression, so as to quantitatively identify over- and under-expressed genes between two or several groups of samples.

While the functionality of these packages goes beyond differential expression analysis, their essential differential expression analysis features are ported to Python in InMoose, respectively in the modules *limma, deseq2* and *edgepy*. Note that *pycombat* relies on functions provided by *edgepy* for RNA-Seq data – following the same dependency as their R counterparts *ComBat-Seq* and *edgeR*.

Despite previous attempts at porting those tools to Python (22), InMoose is, to our knowledge, the sole Python package to provide differential expression analysis for both technologies.

### c. splatter – RNA-Seq and single-cell RNA-Seq Data Simulation

Tool developers often face the challenge of testing their methods against undisputed reference data, to ensure their correctness. Despite its abundance on public databases, real-world data is rarely adequate for this task: its variability cannot be controlled, the required reference annotations may be missing, inconsistent or untrustworthy. Simulated data (also known as synthetic data) provides a way to generate data that *mimics* the behavior of real-world data while controlling its features.

Splatter (5) is a state-of-the-art package to simulate RNA-Seq and single-cell RNA-Seq data in a highly customizable way. In particular, it can model batches and condition groups, features of high interest for batch effect correction and differential expression analysis tools. Splatter can also infer its simulation parameters from real-world data, but this capability has not yet been ported to InMoose.

### d. diffexp – Differential Expression Meta-Analysis

In addition to the three differential expression analysis modules available in InMoose, the module *diffexp* provides a meta-analysis feature to combine the results obtained on different cohorts and/or from different tools. The module relies on a common data structure to store the differential expression results. Meta-analysis is performed on both effect sizes (log-fold-changes) – using a random-effect model – and on *p*-values – using Fisher’s combined probability test and Benjamini-Hochberg adjustment for multiple testing.

While the underlying methods are not novel, harmonizing the format of three state-of-art tools and providing meta-analysis capabilities out-of-the-box streamlines the workflow for bioinformaticians, and seems unique in the ecosystem.

### e. consensus clustering – Cohort Stratification

InMoose also ports an unsupervised clustering algorithm (5), called consensus clustering, that can be used to stratify a cohort. The algorithm identifies clusters in molecular data and provides metrics to help identify the number of clusters that best fits the data. While not exemplified in the present paper, we use it internally to identify sub-cohorts of interest in large cancer cohorts (23).

## Supporting information

Code to reproduce the workflow and the figure

## Acknowledgements

The authors would like to thank Benjamin Millot and Michael I. Love for their support and valuable comments.

