## Supplementary material for "Bulk Transcriptomic Analysis with InMoose, the Integrated Multi-Omic Open-Source Environment in Python": Code to reproduce the workflow and the figure: README.md

This supplementary material contains a single file that contains:
- the code from listing 1
- the code from listing 2
- the code to regenerate the figure

Install the dependencies specified in the `requirements.txt` file:
```
pip install -r requirements.txt
```
then run the code:
```
python workflow.py
```

The figure will then be stored in a file named `pcas.png`.
