## Supplementary material for "Bulk Transcriptomic Analysis with InMoose, the Integrated Multi-Omic Open-Source Environment in Python": Code to reproduce the workflow and the figure: workflow.py

```
import matplotlib
import matplotlib.pyplot as plt
import numpy as np
import pandas as pd
import scanpy as sc
from anndata import AnnData
from inmoose.deseq2 import DESeq, DESeqDataSet
from inmoose.diffexp import meta_de
from inmoose.pycombat import pycombat_seq
from inmoose.sim import sim_rnaseq
from scipy.stats import pearsonr

## LISTING 1 ##

# number of genes
N = 10000
# number of samples
M = 2000
assert M % 10 == 0
P = M // 10  # 10% of M, helper variable

# 3 batches: 20% 30% 50% of the samples
batch = (2 * P) * [0] + (3 * P) * [1] + (5 * P) * [2]
batch = np.array([f"batch{b}" for b in batch])
batch0 = batch == "batch0"
batch1 = batch == "batch1"
batch2 = batch == "batch2"

# 2 condition groups
#   - group 1: 50% batch 1, 33% batch 2, 60% batch 3
#   - group 2: 50% batch 1, 67% batch 2, 40% batch 3
group = P * [0] + P * [1] + P * [0] + (2 * P) * [1] + (2 * P) * [0] + (3 * P) * [1]
group = np.array([f"group{g}" for g in group])
assert len(batch) == M and len(group) == M

# store clinical metadata (i.e. batch and group) as a DataFrame
clinical = pd.DataFrame({"batch": batch, "group": group})
clinical.index = [f"sample{i}" for i in range(M)]

# simulate data
# random_state passes a seed to the PRNG for reproducibility
counts = sim_rnaseq(N, M, batch=batch, group=group, random_state=42).T


## LISTING 2 ##

# run AD meta-analysis on batches 1, 2 and 3
# first run the differential expression analysis on each batch individually
cohorts = [
    DESeqDataSet(counts[b], clinical.loc[b], design="~ group")
    for b in [batch0, batch1, batch2]
]
individual_de = [DESeq(c).results() for c in cohorts]
# then aggregate the results
ad = meta_de([de for de in individual_de])

# run ISD meta-analysis on merged batches
# first correct batch effects to properly merge batches
# transpositions account for the difference of formats for pycombat_seq and deseq2
harmonized_counts = pycombat_seq(counts.T, batch).T
# then run the differential expression analysis on the merged batches
isd = DESeq(DESeqDataSet(harmonized_counts, clinical, design="~ group")).results()


## PLOTS ##

# in a single figure
#   A. PCA before BEC
#   B. PCA after BEC
#   C. heatmap of correlations between results

colors = matplotlib.rcParams["axes.prop_cycle"].by_key()["color"]
pl = {
    "batch0": colors[0],
    "batch1": colors[1],
    "batch2": colors[2],
    "group0": colors[3],
    "group1": colors[4],
}

fig = plt.figure(layout="constrained", figsize=(8, 12))
subfigs = fig.subfigures(3, 1, wspace=0.07, height_ratios=[2, 2, 2])

ax = subfigs[0].subplots(1, 2)
before_bec = AnnData(counts, obs=clinical)
sc.tl.pca(before_bec)
sc.pl.pca(
    before_bec, color="batch", ax=ax[0], show=False, palette=pl, legend_loc="best"
)
sc.pl.pca(
    before_bec, color="group", ax=ax[1], show=False, palette=pl, legend_loc="best"
)

ax = subfigs[1].subplots(1, 2)
after_bec = AnnData(harmonized_counts, obs=clinical)
sc.tl.pca(after_bec)
sc.pl.pca(after_bec, color="batch", ax=ax[0], show=False, palette=pl, legend_loc="best")
sc.pl.pca(after_bec, color="group", ax=ax[1], show=False, palette=pl, legend_loc="best")

elems = ["batch0", "batch1", "batch2", "AD", "ISD"]
hm = pd.DataFrame(index=elems, columns=elems)
ad["log2FoldChange"] = ad["combined logFC"]
ds = {
    "batch0": cohorts[0].results(),
    "batch1": cohorts[1].results(),
    "batch2": cohorts[2].results(),
    "AD": ad,
    "ISD": isd,
}
for e in elems:
    for f in elems:
        corr = pearsonr(ds[e]["log2FoldChange"], ds[f]["log2FoldChange"]).statistic
        hm.loc[e, f] = corr
hm = hm.astype(float)
ax = subfigs[2].subplots(1, 1)
im = ax.imshow(hm, cmap="YlGn")
ax.set_xticks(np.arange(hm.shape[1]), labels=hm.columns)
ax.set_yticks(np.arange(hm.shape[0]), labels=hm.index)

# Let the horizontal axes labeling appear on top.
ax.tick_params(top=True, bottom=False, labeltop=True, labelbottom=False)

# Annotate the heatmap with correlation values
threshold = im.norm(hm.values.max()) / 2.0
for i in range(hm.shape[0]):
    for j in range(hm.shape[1]):
        if im.norm(hm.iloc[i, j]) > threshold:
            cl = "white"
        else:
            cl = "black"
        text = im.axes.text(
            j,
            i,
            f"{100*hm.iloc[i, j]:.2f}%",
            horizontalalignment="center",
            verticalalignment="center",
            color=cl,
        )

subfigs[0].suptitle("A", ha="left", va="top", fontweight="bold", fontsize="xx-large")
subfigs[1].suptitle("B", ha="left", va="top", fontweight="bold", fontsize="xx-large")
subfigs[2].suptitle("C", ha="left", va="top", fontweight="bold", fontsize="xx-large")

fig.savefig("pcas.png")
```
